# High-depth Whole Genome Sequencing of Single Human Colon Crypts Uncovers New View on Crypt Clonality

**DOI:** 10.1101/2025.09.24.677335

**Authors:** Zarko Manojlovic, Cindy Okitsu, Timothy Okitsu, Jordan Wlodarczyk, Michael R. Lieber, Yong Hwee Eddie Loh, Chih-Lin Hsieh

## Abstract

It is generally accepted that each colon crypt is monoclonal and is populated by a single stem cell lineage after early human life. It has been impossible to profile somatic mutations genome-wide because high-depth and high-quality whole genome sequencing (WGS) of single cells is unachievable without tissue culture or whole genome amplification (WGA). Therefore, the cell-to-cell heterogeneity in each individual remains mostly unknown. Applying our novel WGA-free WGS method to obtain >30X post-alignment depth, we comprehensively profiled somatic mutations in 71 single human colon crypts with matched bulk controls from 14 individuals. Analysis reveals that colon crypts are commonly of multiple lineages in human adults. Our study is the first to demonstrate that an appropriately designed WGS approach can determine cell-to-cell heterogeneity in natural cell clones. The much higher sensitivity of WGS than the previous methods in lineage tracing can unlock the complex stem cell dynamics in the colon crypt.

## INTRODUCTION

Human intestinal stem cells (hISCs) are responsible for the constant regeneration of the colon crypt epithelial lining, which is replaced as frequently as every three to five days (Kosinski et al., 2007; Kozar et al., 2013; Nicolson et al., 2018). hISCs reside in a stem cell niche at the base of the crypt and divide to form new stem cells and transit-amplifying (TA) cells. TA cells proliferate, become differentiated after migrating upward, and shed into the colon lumen after reaching the crypt orifice (Clevers, 2013; Kershaw et al., 2013). Based on lineage tracing experiments using single or a few markers, it has been proposed that neutral drift leads to monoclonality of the crypts in mice in 1 to 6 months, with a single ISC giving rise to all the cells in a crypt of a mouse after 3 months (Nicolas et al., 2007; Snippert et at., 2010; Reizel et al., 2011; Clevers, 2013; Clevers and Batlle, 2013; Kozar et al., 2013). Studies using genetic mutations in nuclear and mitochondrial genomes have concluded that there are 5 to 7 ISCs in each human colon crypt to maintain the stem cell niche, and monoclonal conversion of human colon crypts takes 6 to 13 years (Baker et al., 2014; Nicholson et al., 2018; Stamp et al., 2018). Therefore, it has been commonly accepted that each colon crypt, which consists of 1,000 to 2,000 cells, is monoclonal and is populated by a single stem cell lineage after early human life.

The human somatic cell genome becomes increasingly heterogeneous as each person develops and ages over time due to acquired DNA mutations caused by DNA damage and replication errors. Although somatic heterogeneity within an individual is not heritable across generations, the impact of acquired mutations in somatic cells is apparent in cancer, degenerative diseases, and mitochondrial diseases. Heterogeneity or genetic variation within the individual has been difficult to address, and the extent of cell-to-cell variation within each individual has been quite unclear. It has been impossible to obtain high-depth and high-quality genome-wide sequences from individual cells without tissue culture or extensive whole genome amplification (WGA), which can introduce DNA mutations and genome coverage bias. These problems persist despite the recent development of single-cell sequencing technology (Ruan et al., 2020; Biezuner et al., 2021; Fan et al., 2021; Gonzalez-Pena et al., 2021).

We modified the conventional whole genome sequencing (WGS) library preparation method to generate balanced ≥30X depth post-alignment high-quality genomic sequences from 14 individuals (71 single colon crypts and 14 bulk DNA controls) without WGA or DNA purification (Manojlovic et al., 2023). Our study was designed to identify somatic DNA mutations from each single colon crypt to reveal the accumulation of mutations and the increase of heterogeneity with age. Somatic DNA mutations identified in these single crypts would represent mutations in the stem cells that give rise to the crypt lining and mutations that occur during the first few proliferative divisions of the TA cells that populate the crypt. With the assumption that each colon crypt is monoclonal after the early stage of human life, colon crypts can be used as a model micro-organ system to identify mutations occurring in single stem cells and reveal the extent of cell-to-cell variation within each individual to understand somatic heterogeneity. Our high-depth WGS dataset generated from natural cell clones can provide a comprehensive profile of mutations in each colon crypt and new insights into hISC biology.

If each colon crypt were monoclonal, as inferred from previous stem cell lineage tracing studies using single genetic markers, then mutations from the original single stem cell progenitor should be present in all the cells in the crypt. Somatic autosomal mutation in the stem cell progenitor that occurs only on a single chromosome in the diploid cell should have a variant allele frequency (VAF) of 0.5 in a monoclonal crypt cell population. Stem cells accumulate new replication independent mutations while residing in the stem cell niche of each colon crypt, and replication-dependent mutations arise in each daughter cell during cell division to repopulate the stem cell niche and form the crypt lining. These new mutations would be present at VAFs lower than 0.5 in the crypt cell population, depending on the number and lineage of the stem cells participating in the crypt lining formation (assuming no elevated cell death or acquired proliferative advantage in the daughter cells). The VAF distribution profile of autosomal variants in a single stem cell lineage crypt would have a major binomial distribution peak at VAF 0.5 and some other minor distribution peaks at VAF lower than 0.5 depending on the length of time and the branching points for a single lineage to populate the niche. In contrast, the distribution peak of the highest VAF would be at a VAF <0.5 if stem cells from more than a single lineage participate in the crypt lining formation. Therefore, the VAF distribution profile of each crypt allows the determination of whether a crypt is formed by single or multiple stem cell lineages. Similarly, autosomal variants shared by all colon crypts sequenced from the same individual should have a VAF of 0.5 since these variants must exist in their common stem cell progenitor before these crypts become independent micro-organs. A conclusion that crypts were derived from a single ancestral stem cell was presented in a previous low-depth WGS study (Lee-Six et al., 2019).

While it is commonly accepted that each colon crypt is formed by a single stem cell lineage after early human life, it remains controversial whether homeostatic maintenance of the stem cell niche is by symmetrical or asymmetrical division (Lopez-Garcia et al., 2010; Snippert et al., 2010; Reizel et al., 2011; Stamp et al., 2018). After each division, half of the daughter cells from stem cells in the niche remain in the niche for homeostatic maintenance and the other half migrate out to become TA cells forming the crypt lining. If stem cells in the niche divide symmetrically, daughter cells of all stem cells have equal opportunity to remain in the niche or to migrate out after each cell division. With a symmetrical division mechanism, the genetic makeup in the niche would be dynamic with each generation of stem cell division and neutral competition. If stem cells in the niche divide asymmetrically, one of the two daughter cells from each stem cell will remain in the niche while the other one exits the niche after each cell division in general. If asymmetrical division were the mechanism for niche maintenance, each crypt would have the same composition of stem cells in the niche from the time a crypt is derived in early development, and the genetic makeup of each crypt would not change dramatically over time. Therefore, it would take much longer for each crypt to reach a monoclonal state if stem cells divide asymmetrically than if they divide symmetrically. It has been suggested that allele fractions (frequencies) of DNA mutations from individual crypts may clarify this debate (Stamp et al., 2018).

Our single colon crypt study reported here is the first to achieve high-depth and high-quality WGS on uncultured single human natural cell clones with a matched germline control from each individual. The high sensitivity of WGS in lineage tracing reveals the complex dynamics of stem cell activity in the colon crypts and shows that colon crypts are commonly polyclonal in adult humans. WGS can identify somatic mutations reliably and comprehensively to provide new evidence at high resolution for colon crypt clonality and potentially, the stem cell self-renewal mechanism.

## MATERIALS AND METHODS

### Tissue collection and crypt isolation

Colon samples were collected from 14 individuals ages 20 to 90 years old who had undergone surgery to remove part of the colon as a step in the standard of care for their diseases at Keck Hospital of the University of Southern California through the Translational Pathology Core with an approved IRB. An approximately 5 mm x 5 mm colon specimen is used to isolate 20 to 30 individual crypts under an inverted microscope as previously described (Manojlovic et al., 2023). The crypts are stored at -80°C after confirming that only a single colon crypt is present in each tube under the microscope. The remaining crypts in the suspension are spun down for bulk DNA extraction, and the extracted DNA is stored at 4°C.

### Whole genome sequencing of colon crypt and bulk tissue libraries

Sequencing libraries from single colon crypts and the bulk DNA are constructed using the NEBNext Ultra DNA Library Prep Kit (NEB #E7370), as described in detail previously (Manojlovic et al., 2023). After quality and quantity assessment of each sequencing library, the libraries are pooled for 150 bp paired-end sequencing on S4v1.5 flow cells using NovaSeq 6000 (Illumina, San Diego). The post-alignment statistics of 91% > 20x and quality measures of these libraries are provided in a previous publication (Manojlovic et al., 2023).

### Variant Calling

FASTQ reads were aligned to GRCh38 using BWA-MEM v0.7.17.

Duplicate reads were marked with Picard, and base quality recalibration was performed using GATK v4.2 Best Practices by the USC Keck Genomics Platform (KGP) in-house pipeline as described previously (Manojlovic et al., 2023). Somatic SNVs were called using two independent algorithms, Mutect2 (GATK) and Strelka v2.9.0, using matched bulk colon DNA as the germline control to reduce germline variant contamination. Two germline callers, HaloptypeCaller and Freebayes v1.2.0, are used to generate germline variant calls. Combining results from different callers was suggested to obtain more reliable variant calls because different somatic variant caller algorithms have their own strengths and weaknesses (Cai et al., 2016). SNVs were retained only if called by both Mutect2 and Strelka (2Caller consensus). Our 2Caller consensus algorithm is described in detail and available on GitHub (https://github.com/twewyttst777/merge_vcf).

In addition, we performed an independent commercially available pipeline, DRAGEN Somatic v4.2.4 (Illumina Inc.), to validate caller performance on all samples. We concluded that the two pipelines have a high inter-caller concordance with 93.78% (s.d. 3.10%) 2Caller calls present in DRAGEN and 80.72% (s.d. 7.91%) DRAGEN calls present in 2Caller (Supplementary Table S1). The variant calls from 2Caller are used to increase variant call confidence for all downstream analyses. The variant calls from Illumina’s DRAGEN somatic pipeline are not used in the downstream analysis of our data, other than filtering for read depth for specific analysis as indicated and for internal validation of the analysis.

### Generation of VAF distribution profile

The reliability of single-nucleotide variant (SNV) identification is much higher than that of insertion/deletion analysis. Also, males and females have different sex chromosome complements, which would affect the allele frequency calculation of variants on those chromosomes. In addition, the reliability of VAF increases with higher read depth at the variant position. Therefore, only autosomal SNV with a total read depth ≥15X is included in this analysis and the X chromosome SNVs are analyzed separately without depth filtering due to the lower read count in males. The VAF is calculated by dividing the alternate (mutant) allele read depth by the total read depth at the variant position. Two VAF distribution profiles, one for autosomes and one for the X chromosome, were generated using Metplotlib (Hunter, 2007) with Python v3.6.9. The graph is presented by binning VAF (0.01 bins for autosomes and 0.02 bins for the X chromosome) and plotting the count of variants in each bin from VAF 0 to 1.0. The 0.02 bin is used for the X chromosome due to the lower counts of variants on the X chromosome, which accounts for about 8% of the human genome, compared to all autosomes.

### Obtaining and reanalyzing previously published low-depth WGS data

A WGS dataset including 2038 CRAM files of sequencing reads from 440 crypts (42 individuals, as listed in Supplementary Tables 1 and 2 in Lee-Six et al. 2019) was downloaded with a signed data transfer agreement (called Sanger study here). The majority of the 440 samples have multiple CRAM files for each sample, which required performing Samtools v1.19.2 merge before running Illumina’s Dragen Somatic v4.2.4 pipeline in Tumor-Only mode. We tested the tools for merging the multiple CRAM files of the same sample to ensure that no data is lost in the process. A total of 28 crypts with the highest median read depth >15X from four individuals were chosen for analysis (see Supplementary Methods for details and rationale).

A total of 86 CRAM files from 28 crypts were processed through the DRAGEN pipeline (Supplementary Table S2), and only the SNV calls, which are known to be the most reliable variant calls from WGS analysis pipelines, were included in the downstream analysis (Supplementary Methods). As described above, the VAF of autosomal SNVs are calculated to generate the VAF distribution profile for each crypt. The VAF distribution profile of SNVs from each crypt is plotted with Excel after each step of the two-step (dbSNP removal and shared variant removal) germline variant contamination reduction since no matched-germline control was in the Sanger study. A VAF distribution profile of shared variants removed in the second step of the germline variant removal strategy is also plotted for each crypt to demonstrate the germline nature of these removed variants from the dataset.

## RESULTS

### Expected autosomal VAF distribution profile representing colon crypts of single lineage or multiple lineages

If multiple stem cells from a single lineage populate each crypt (after 6 to 13 years in humans), then the autosomal VAF distribution profile in each adult crypt is expected to show a binomial distribution around the VAF peak at 0.5. This distribution represents somatic variants existing in all the stem cells, which derived from a single progenitor. It is important to note that somatic variant callers identify new variants arising in each colon crypt that are absent in the bulk tissue. Thus, the VAF analysis does not include germline variants present in all cells of the individual. Additional binomial distribution peaks, if they reach a level of detectability, would have VAFs <0.5 and represent somatic variants arising in the stem cell self-renewal and TA cell proliferation divisions.

For a stem cell lineage to fully populate the stem cell niche, both daughter cells from multiple generations of the lineage progenitor stem cell must remain in the niche if 5 to 7 stem cells are in the niche for homeostatic maintenance. If asymmetrical division (only one daughter cell from each stem cell remains in the niche after each cell division) is the primary mechanism for homeostatic maintenance in the niche, it may take a much longer time for a crypt to become a single stem cell lineage than if stem cell division is symmetrical (both daughter cells may remain in the niche after each stem cell division).

New and different mutation(s) continue to arise in each daughter cell and its descendants with each round of replication after the two daughter cells branch from the same progenitor. Depending on the length of time stem cells from the same lineage fully populate the stem cell niche, these stem cells can harbor many different DNA mutations accumulated through DNA replication-dependent and replication-independent mechanisms despite originating from the same progenitor. The height and location of the VAF distribution peaks of these new mutations would depend on the number of stem cells participating in the crypt lining formation, how many generations the lineage progenitor took to fully populate the niche, and the branching point of the multiple stem cells from the lineage. If a crypt is fully populated by stem cells from a single lineage within a few successive cell divisions, the VAF distribution profile of the crypt would be as if it contains only a single stem cell. If it takes years for a crypt to become fully populated by stem cells from a single lineage, the crypt would have additional peaks at VAF <0.5 beyond the peak at VAF 0.5 in the VAF distribution profile. If the crypt lining is formed from stem cells of different lineages, the VAF distribution peak at 0.5 may be entirely absent. The VAF distribution profile derived from the high-depth WGS would be sufficiently sensitive to reflect whether the crypt lining is formed from stem cells of a single lineage or multiple lineages.

### Autosomal VAF distribution profile reveals stem cell clonality and the dynamic stem cell activity in crypt lining formation

Three distinct VAF distribution profiles, A, B, and C, of autosomal SNV are observed among the 71 single colon crypts from 14 individuals of 20 to 90 years old in our dataset (Supplementary Table S1 and Supplementary Figure S1 left panels for all 71 crypts). These three VAF distribution profiles are described with a representative example for each below.

Profile A, which represents a single stem cell lineage crypt, has a major VAF distribution peak at ∼ 0.5 as described above (Figure 1A). Profile B, which represents a crypt of multiple lineages (multi-lineage), typically has a major distribution peak at VAF ≤0.25 without a distribution peak at VAF 0.5, indicating the lack of shared variants (which have a theoretical VAF 0.5) beyond the variants existing in the bulk tissue of the colon (Figure 1B). Profile C has distribution peaks at varied VAFs in addition to a peak at VAF 0.5 (Figure 1C). The peak at VAF 0.5 indicates that all stem cells involved in forming the crypt have a common progenitor and any additional peaks between VAF 0.5 and VAF 0.25 represent variants present only in a fraction of the stem cells. Profile C represents a crypt populated by stem cells that diverged from a common progenitor at far-apart lineage branching points and should be considered as different stem cell lineages on the molecular level. The presence of extra or shifted VAF peaks in Profiles B and C depends on the number of stem cells, their lineages, and the proportions of each lineage contributing to the specific crypt formation.

**Figure 1.**
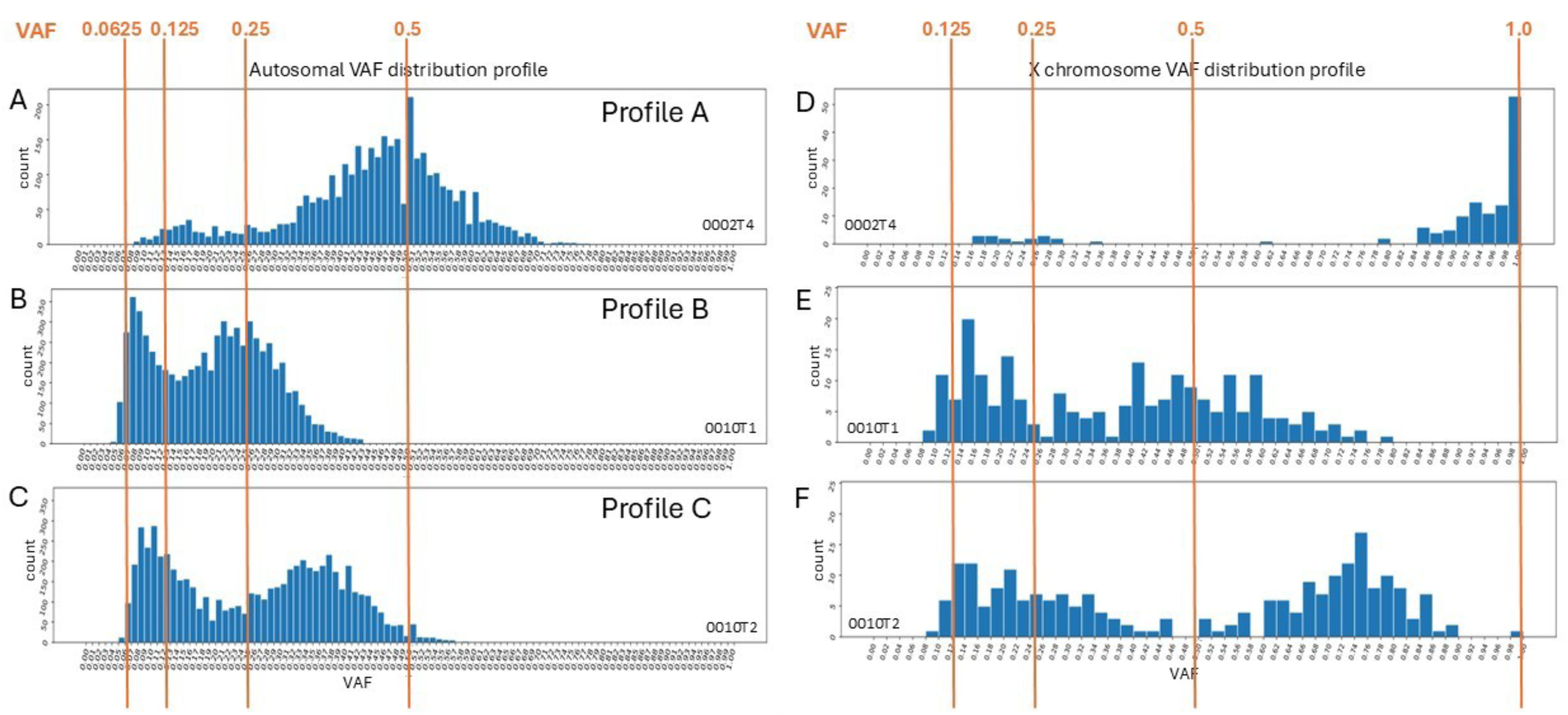
Representative VAF distribution profiles of autosomal and X chromosome single nucleotide variants for single human colon crypts. Autosomal single nucleotide variant (SNV) calls with a total read count ≥15 from our 2caller pipeline (common calls from Mutect and Strelka) are included in this analysis. VAF is calculated by dividing the alternate (mutant) allele read count by the total read count at the variant position. A) Autosomal VAF distribution profile of a crypt with a single stem cell lineage. B) Autosomal VAF distribution profile of a crypt with multiple stem cell lineages (no shared variants among the lineages). C) Autosomal VAF distribution profile of a crypt with multiple stem cell lineages with a distant common origin. D), E), and F) are X chromosome VAF distribution profiles of the same crypts presented in A), B), and C), respectively. The VAF of the autosomal variants are binned by 0.01 in plots A, B, and C. The VAF of the X chromosome variants are binned by 0.02 (due to the lower count of X chromosome variants than autosomal variants) in plots D, E, and F. The X-axis is the VAF and the Y-axis is the variant count. Please see Supplementary Figure S1 for VAF profiles of all 71 crypts.

Among the autosomal VAF distribution profiles of the 71 single colon crypts sequenced, 28 (39.4%) are categorized as Profile A, 30 (42.3%) as Profile B, and 13 (18.3%) as Profile C (Supplementary Table S1). The previous studies using a single genetic marker for lineage tracing categorized the crypts as marker negative (uninformative for lineage), partially marker positive (multi-lineage), and fully marker positive (single lineage). The feasibility and economics of performing high-resolution WGS increase the sensitivity necessary to confidently categorize the nature of stem cell lineage in colon crypts, exceeding the limitations of previous methods. WGS can categorize every crypt, including the crypts that would have been considered uninformative by the previous studies. Furthermore, the fully marker positive crypts categorized by the previous studies can be divided into Profile A (stem cells fully populate the crypt within a relatively short period of time after branching off the single progenitor) and Profile C (stem cells diverged from a single progenitor at distant time points that should be considered as multi-lineage molecularly). About 58% (Profiles A and C) of the 71 crypts sequenced in our study could have appeared to be single lineage if a single marker were used for lineage tracing, as in previous studies mentioned above. It is important to note that all cells continue to diverge with the accumulation of new replication-dependent and replication-independent mutations in each cell over the human lifespan. Therefore, the stem cells in the niche can become sufficiently diverged to be viewed as different lineages even though they had a common progenitor in the distant past. Instead of being viewed as the end state of monoclonal conversion, Profile A crypts may be considered being in a transient monoclonal state of the dynamic stem cell activity in the niche.

WGS of single crypts, which represent a natural clonal cell population, allows the detection of somatic changes in the human genome at a single base pair level when such information cannot be obtained reliably by WGS of a population of somatic cells.

Our study demonstrates that high-depth and high-quality WGS with matched germline control can reveal the state of molecular heterogeneity in the colon crypt stem cells.

Our analysis indicates that the stem cell activity in populating the stem cell niche and forming the crypt lining may be more complex and dynamic than inferred from previous studies of limited sensitivity. It is possible that the stem cell activity in the crypt is a dynamic process with more than one cycle of convergent and divergent states due to the continuous development of new lineage branching points and the acquisition of new mutations over time.

### VAF distribution profile of X chromosome variants supports the interpretation of the autosomal VAF distribution profile in males

Human males only have a single copy of the X chromosome; therefore, the somatic variants on the X chromosome present in all the stem cells should have a VAF of 1.0 if the crypt lining is derived from a single stem cell lineage. The VAF of X chromosome somatic variants is a powerful method to confirm whether a single or multiple stem cell lineages contribute to the crypt lining formation in males. The corresponding VAF plot of somatic SNVs on the X chromosome was generated for each crypt from all 14 individuals (Supplementary Figure S1 right panels for all 71 crypts).

Crypts with autosomal VAF distribution Profile A showed a one-sided VAF distribution peak at VAF 1.0 for the X chromosome, as expected for a crypt of single stem cell lineage (Figure 1D). Crypts with autosomal VAF distribution Profile B showed multiple VAF distribution peaks without any variants at VAF 1.0 for the X chromosome, consistent with multiple stem cell lineages in the crypt (Figure 1E). Crypts with autosomal VAF distribution Profile C have variants of VAF 1.0 without a typical one-sided binomial distribution for the X chromosome in addition to the multiple VAF distribution peaks (Figure 1F). While the low number of X chromosome variants (X chromosome only accounts for about 8% of the human genome) limits the confidence in categorizing the crypts if considered alone, the X chromosome VAF distribution profile nevertheless provides additional confirmation in males that multi-lineage crypts are common.

### Allele frequency distribution profile of SNVs shared by all crypts from the same individual supports the finding of multi-stem cell lineage crypts in adults

Autosomal somatic variants present in all the crypts sequenced from the same individual must have arisen before these crypts developed from their common progenitor; therefore, such variants should have a theoretical VAF of 0.5 in the crypt cell population if each crypt is of a single stem cell lineage. The autosomal SNV shared by all crypts from the same individual should have a single VAF distribution peak at 0.5 VAF, representing the variants present in the progenitor of all the crypts. However, if a crypt is populated by stem cells of multi-lineage, the VAF distribution for variants shared by all crypts can deviate from the normal distribution peak of 0.5, depending on the representation of this shared lineage in each crypt.

A total of 110 unique variants (distinct from other individuals) are shared by all crypts sequenced from the same individual (12 individuals with 5 crypts sequenced, 1 individual with 6 crypts sequenced, and one individual has no shared variants in 5 crypts sequenced) in our study (Figure 2A). The VAF distribution profile of these 110 unique SNVs (556 total counts) showed a peak around VAF 0.46 with a very wide distribution range instead of the expected VAF 0.5 (Figure 2B). The skewed VAF peak toward lower VAF indicates the presence of additional stem cell lineages that reduce the VAF of the shared variants in at least some crypts from the same individual. This finding supports the conclusion that it is not uncommon for stem cells of multi-lineage to form the lining of a single crypt in human adults.

**Figure 2.**
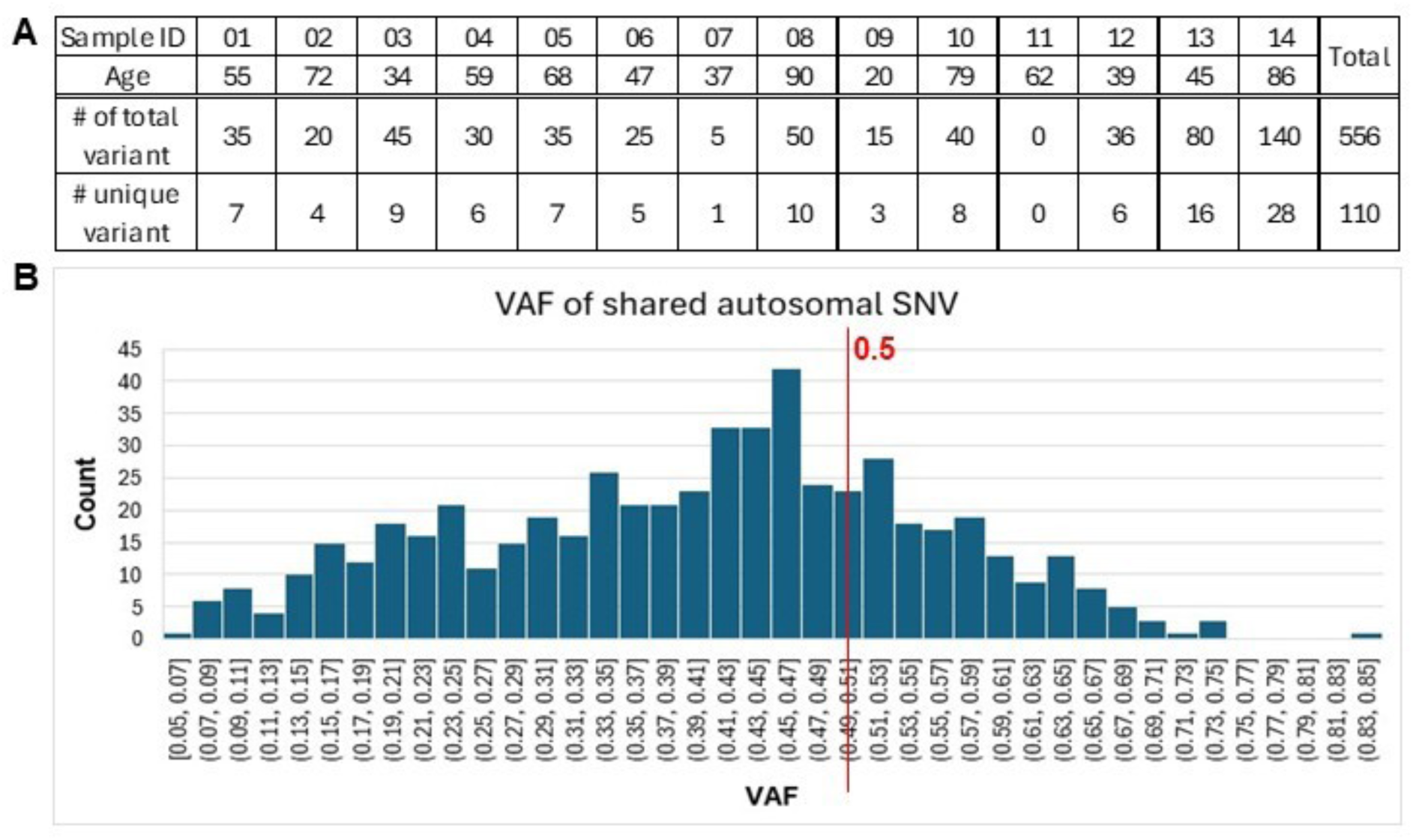
VAF distribution profile of autosomal SNVs shared by all crypts within each individual. A) A total of 556 autosomal SNV calls shared by all crypts from the same individual are included in this analysis. B) VAF distribution of shared variants by all the crypts from the same individual. The Y-axis is the variant count in each bin of 0.02 from VAF 0.05 to 0.85 on the X-axis. Although many variants shared by all crypts sequenced from the same individual have a VAF near 0.5, some are lower than 0.1. A skewed peak centered around lower VAFs (rather than the expected 0.5) deviates from the expected pattern of monoclonal crypts.

### Reanalysis of published low-depth WGS data supports the common presence of multi-lineage crypt in human adults

A published low-depth WGS study of 571 microdissected human colon crypts without matched germline control made the conclusion that all colon crypts are of a single ancestral stem cell (Lee-Six et al., 2019). This conclusion was based on a median VAF of each crypt in a distribution plot. However, the median VAF of each crypt alone is insufficient for determining whether each crypt is of single or multiple lineages, especially given the absence of a matched germline control for each individual (Supplementary Methods). After obtaining the dataset containing 440 crypts from the Sanger Institute (U.K.), CRAM files of 28 WGS crypt samples across four individuals from the Sanger study were processed through the DRAGEN pipeline for variant identification (for basic information on the crypts, see Supplementary Table S2). VAF distribution profiles for these 28 crypts were generated after reducing germline variant contamination in the dataset by a two-step strategy (Supplementary Methods).

Variants in the dbSNP were removed as the first step of the germline variant reduction strategy and the results of a representative crypt are presented and described as follows. The asymmetry of the distribution around the peak at VAF 0.5 is clearly visible in some of the crypts after dbSNP removal (Figure 3 row A, marked by a red arrowhead and Supplementary Figure S2 row A for all 28 crypts). However, potential deviation of VAF distribution profiles from some crypts is not as easily discernable after dbSNP removal (Figure 3, row A, marked by a red question mark and Supplementary Figure S2, row A for all 28 crypts).

**Figure 3.**
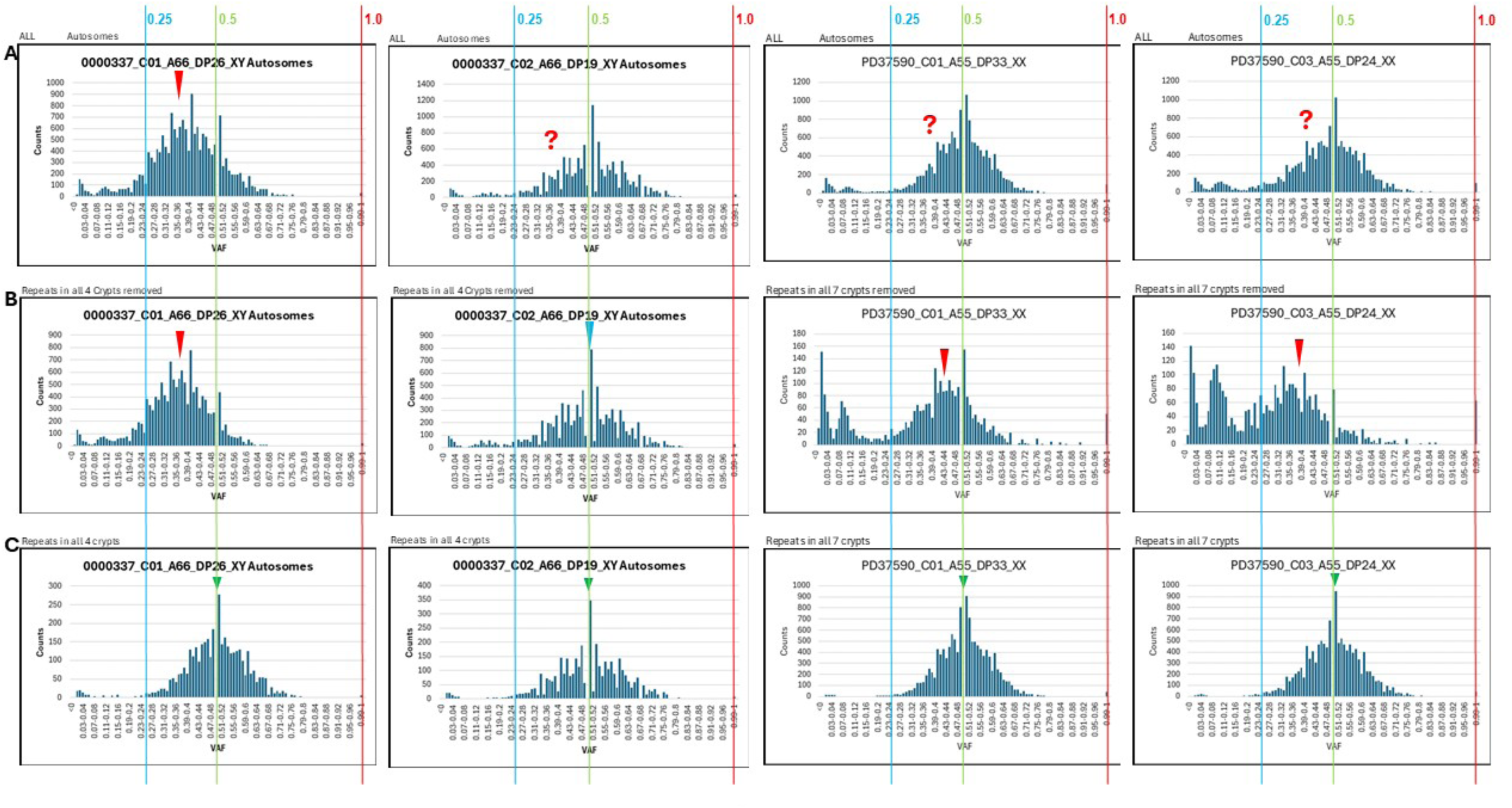
VAF distribution profile of four representative crypts from the Sanger study. The two columns on the left side are two crypts from individual O337 and the two columns on the right side are two crypts from individual PD37590. Row **A)** shows the VAF distribution profile of all the autosomal SNV after removing the dbSNP variants as the first step to reduce germline variant contamination. Row **B)** shows the VAF distribution profile of autosomal variants after the second germline variant contamination reduction step of removing variants shared by all the crypts analyzed from the same individual (repeats in all crypts of the same individual removed). The VAF distribution profiles shown in row B are the best approximation of somatic variants in the Sanger study, in which a matched germline control for each individual is not available (see Supplementary Methods for details). Row **C)** shows the VAF distribution profile of the shared variants that were removed (repeats in all crypts of the same individual) in the second step of the germline variant contamination reduction. <u>Deviation of the profile in row B from the profile in row C for the same crypt indicates the multi-lineage nature of the crypt after germline variant reduction.</u> Although germline variant contamination cannot be entirely removed from the crypts in the Sanger study, the shifted or the extra distribution peak (profiles in row B) is clearly evident in three of the four crypts. Crypt 02 from individual O337 has nearly identical VAF distribution in rows B and C with binomial distribution peaks at VAF 0.5 (as marked by a small green arrowhead), indicating that this crypt is of single stem cell lineage. VAF 0.25, 0.5, and 1.0 are marked by blue, green, and red line, respectively as indicated. The red arrowhead in rows A and B marks the distribution peak deviated from VAF 0.5. The red question mark in row A indicates the potential deviation of the distribution peak. The blue arrowhead in row B marks the distribution peak at VAF 0.5 observed in a single crypt after germline variant removal. The individual ID, crypt ID, individual age, median read depth of the sequencing data, and the sex chromosome complement as displayed with an underscore in between are indicated at the top of each profile. The Y-axis represents the count in each bin of 0.01 displayed from 0 to 1 as variant allele frequency on the X- axis. Please see Supplementary Figure S2 for VAF distribution profiles of all 28 crypts from the Sanger study.

Variants shared by all the crypts from the same individual were subsequently removed in the second step of the germline variant reduction strategy (Supplementary Methods). A shifted or extra distribution peak in the VAF distribution profile of three of the four crypts can be clearly discerned after shared variant removal (Figure 3, row B, peaks denoted by the red arrowheads). Only one of the crypts presented here had a VAF distribution Profile A as a single lineage crypt after shared variant removal (Figure 3 row B, blue arrowhead marked the symmetrical distribution peak at VAF 0.5). The VAF distribution profile of the variants removed from each crypt that are shared by all crypts of the same individual showed a symmetrical distribution peaked at VAF 0.5 (peak Figure 3 row C, marked by small green arrowhead and Supplementary Figure S2 row C for all 28 crypts). The high count (Supplementary Table S2) and the pattern of VAF distribution (Supplementary Figure S2 row C) of shared variants by all crypts from the same individual in each crypt indicate the germline nature of these variants.

After removing the potential germline variants, shifted or extra VAF distribution peaks between VAF 0.25 and 0.5 are clearly visible in nearly all 28 crypts from four individuals in the Sanger study (Supplementary Figure S2 row B) even though there are an easily appreciable number of variants with VAF 1.0, which indicates the incomplete germline variant removal, remaining in the dataset. A total of 730 variants (average 0.88%, s.d. 0.53%) of VAF 1.0 remained in 102,408 total variants after the 2-step germline variant removal in the 28 crypts from the Sanger study (Supplementary Table S2).

In contrast, the somatic variant calls from the 71 single colon crypts in our dataset were made by referencing a matched bulk tissue control from the same individuals. The very low likelihood of germline contamination in our dataset is demonstrated by only 28 of the 307,229 total variants in 71 crypts (<0.01%) are of VAF 1.0 (Supplementary Table S1). Without a germline control, the overwhelming number of contaminating germline variants can conceal the VAF distribution peak of somatic variants and lead to erroneous interpretation of the results. However, over-removal of somatic variants does not affect the VAF distribution profile (Supplementary Methods). This reanalysis exercise highlights the difficulty in the complete removal of germline variant contamination if a control sample (preferably a tissue-matched control) from the same individual is absent for WGS analysis. Our reanalysis of the 28 crypts from four individuals in the Sanger study clearly showed that human colon crypts are predominantly polyclonal at the molecular level despite the incomplete removal of germline variants from the dataset. This reanalysis also clearly demonstrated that the complete removal of germline variants in the VAF analysis is essential for accurately categorizing the stem cell lineage in colon crypts.

## DISCUSSION

Based on the findings from multiple previous elegant studies using single or a few markers to trace stem cell lineage in colon crypts, it is widely accepted that the stem cell niche at the bottom of each colon crypt is populated by multiple stem cells. With neutral drift and niche succession, it has been proposed that each crypt becomes monoclonal with multiple stem cells of a single lineage populating the stem cell niche after early life in mice and humans. Our study of 71 single human colon crypts with matched bulk tissue controls from 14 individuals ranging from 20 to 90 years of age is the first study to sequence natural cell clones from humans without *ex vivo* culture or WGA to generate high-depth and high-quality WGS data. Our unique WGS method enables a comprehensive analysis of genome-wide DNA mutations in natural cell clones such as single human colon crypts. In our analysis, we unexpectedly discovered that the colon crypts are commonly formed by multiple stem cell lineages in human adults, departing from the previously accepted view of monoclonality.

High read depth WGA is sufficiently sensitive to detect DNA changes at the base pair level with confidence and to provide a genome-wide view of heterogeneity in the stem cells contributing to the crypt lining formation. The VAF distribution analysis allows the distinction of single lineage and multi-lineage crypts and provides further information on the colon stem cell dynamics in the stem cell niche. We categorized the clonality of the stem cells in the colon crypts into three groups: single lineage (Profile A), multi- lineage without common progenitor (Profile B), and multi-lineage with a common distant progenitor (Profile C), based on the distribution peaks observed in the VAF distribution profile of each crypt. Depending on the number of stem cells in the niche and the timing of the branching point, the VAF distribution peaks of the multi-lineage crypt deviate from that of the single lineage crypt. It is important to note that there are potential grey areas in categorizing these VAF distribution profiles in our study because the random sampling effect of molecules in sequencing remains at the >30X high read depth of WGS (WGS of even higher depth may reduce this problem). Nevertheless, it is quite clear that multi-lineage crypts are not uncommon in human adults.

Nearly 40% of the 71 single colon crypts in our study is of a single lineage, with multi-linage crypts accounting for the remaining 60%. This observation differs from the previous conclusion made in a low-read depth WGS Sanger study of human colon crypts (Lee-Six et al., 2019). Our reanalysis, which was necessary due to the lack of detailed information in the previous publication, of data on a subset of 28 colon crypts with the best WGS coverage in the Sanger study clearly shows that the majority of these crypts are also of multi-lineage. This reanalysis unequivocally shows that data from the Sanger study is consistent with and supports our current observation that multi- lineage crypts are common in human adults. Furthermore, the VAF distribution is more likely to show clearer deviation from the single lineage VAF distribution profile than demonstrated in the current reanalysis if the germline variant contamination can be removed entirely from each crypt in the Sanger study. Together, findings in the 71 single crypts from our study and in the subset of 28 crypts from the Sanger study reanalyzed clearly depart from the previously widely accepted view that all crypts are of single lineage after early life in humans.

Importantly, our findings do not contradict the findings of previous studies mentioned earlier using single or a few markers for lineage tracing. In fact, ∼58% of the 71 crypts in our study could have been classified as single lineage if a single DNA mutation were used as the lineage marker. WGS has the resolution to identify all somatic mutations (thousands instead of fewer than a dozen in each crypt) from all stem cell lineages in every crypt including the ones that would be called uninformative by the previous marker-based lineage tracing studies. Therefore, previous marker-based lineage tracing studies likely underestimated polyclonality due to limited resolution and high rates of uninformative crypts. Significantly, our study illustrates the high sensitivity and capability of WGS for stem cell lineage tracing in natural cell clones such as colon crypts and the potential to uncover stem cell activity and dynamics in colon crypts.

Although our study was not designed for stem cell lineage tracing, our study reveals the complexity of stem cell dynamics in the colon crypt. Our experience also strongly suggests that higher sequencing depth is essential for higher accuracy of VAF. In addition, our report here also illustrates that a matched germline control from the same individual, preferably from the same tissue type, is essential for somatic variant discovery and should not be omitted in somatic mutation analysis. WGS studies designed appropriately with adequate statistical power and an appropriate germline variant control will be needed to determine the number of stem cells and the frequency of multi-lineage crypt presence at different ages and to learn the dynamics of stem cell activity in each crypt and how the dynamics change over the human lifespan. If asymmetrical division is the primary mechanism for stem cell renewal in the crypt, the succeeding TA pools would be much more similar than if symmetrical division is the primary mechanism as mentioned earlier. With the sensitivity of high-depth and high-quality WGS, mutations in the upper and lower halves of single colon crypts may be identified to determine whether the stem cell maintenance mechanism is symmetrical or asymmetrical division.

## ACKNOWLEDGEMENT

We would like to thank the Norris Comprehensive Cancer Center Translational Pathology Core for the sample collection. We would like to acknowledge the Norris Comprehensive Cancer Center Molecular Genomics Core and the Keck Genomics Platform at the University of Southern California for the sequencing work. This work is supported by funds from NIA R01 AG 067615 and the Catherine and Joseph Aresty Endowment.

---

**Supplementary Figure S1.**
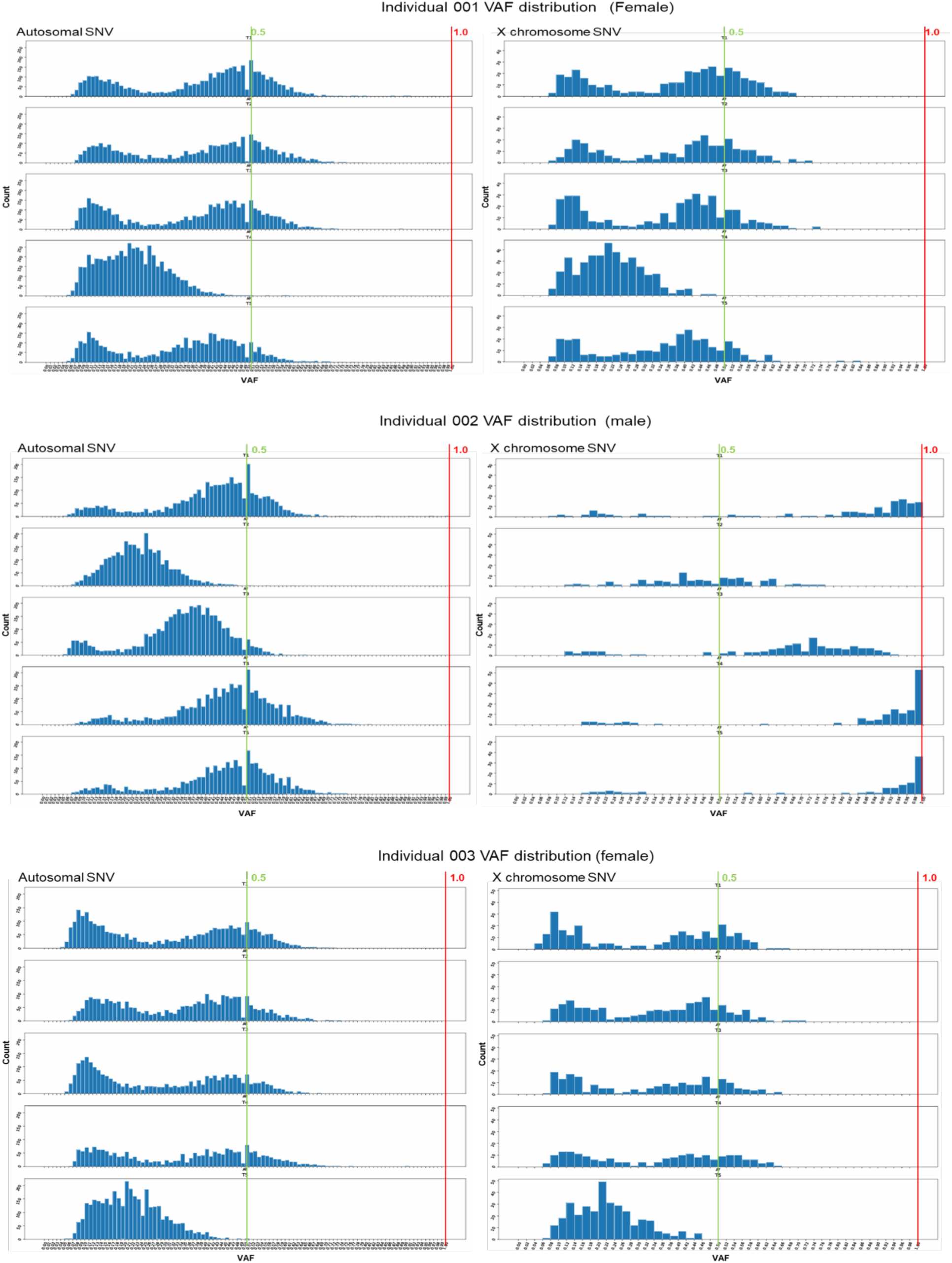
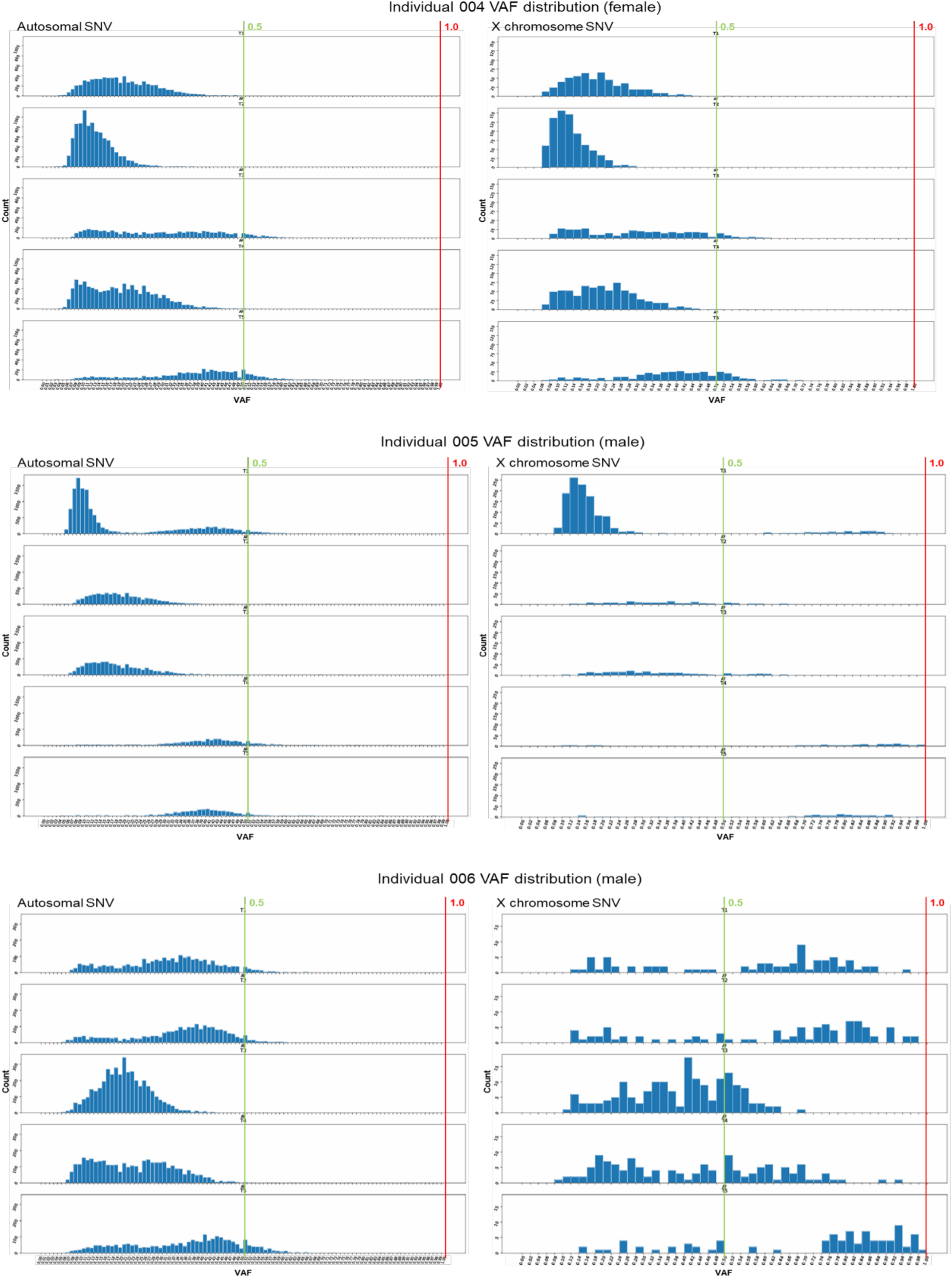
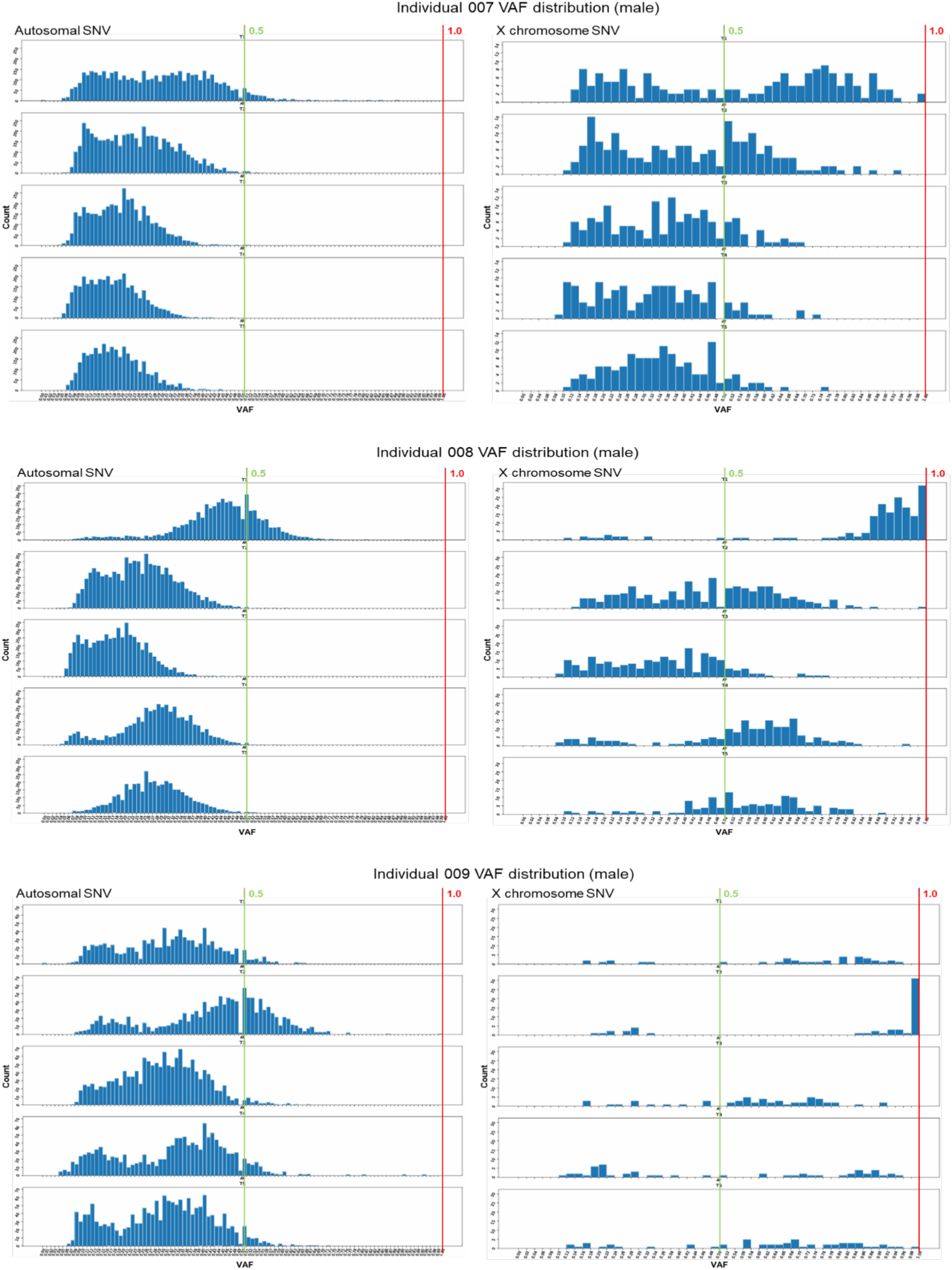
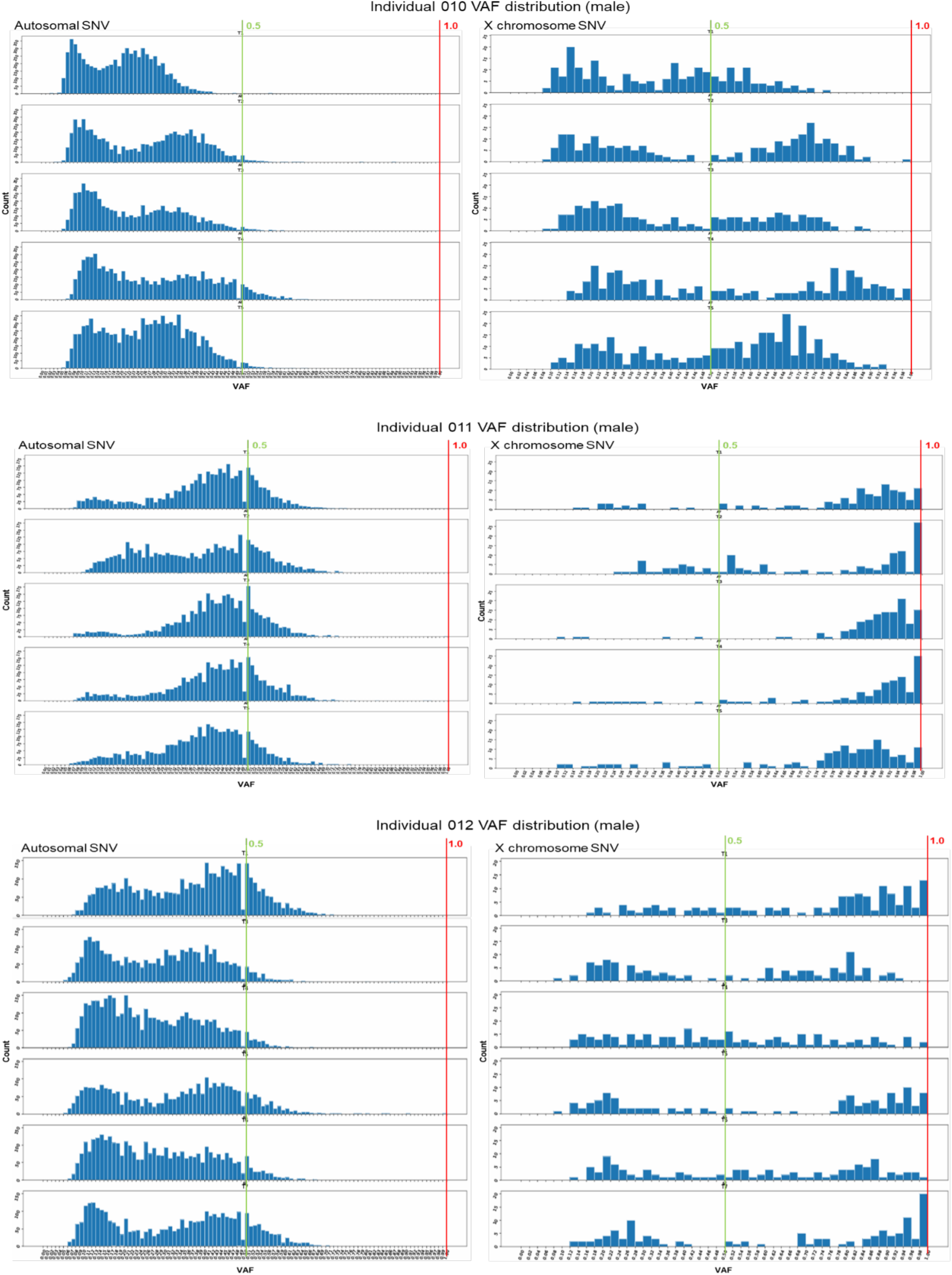
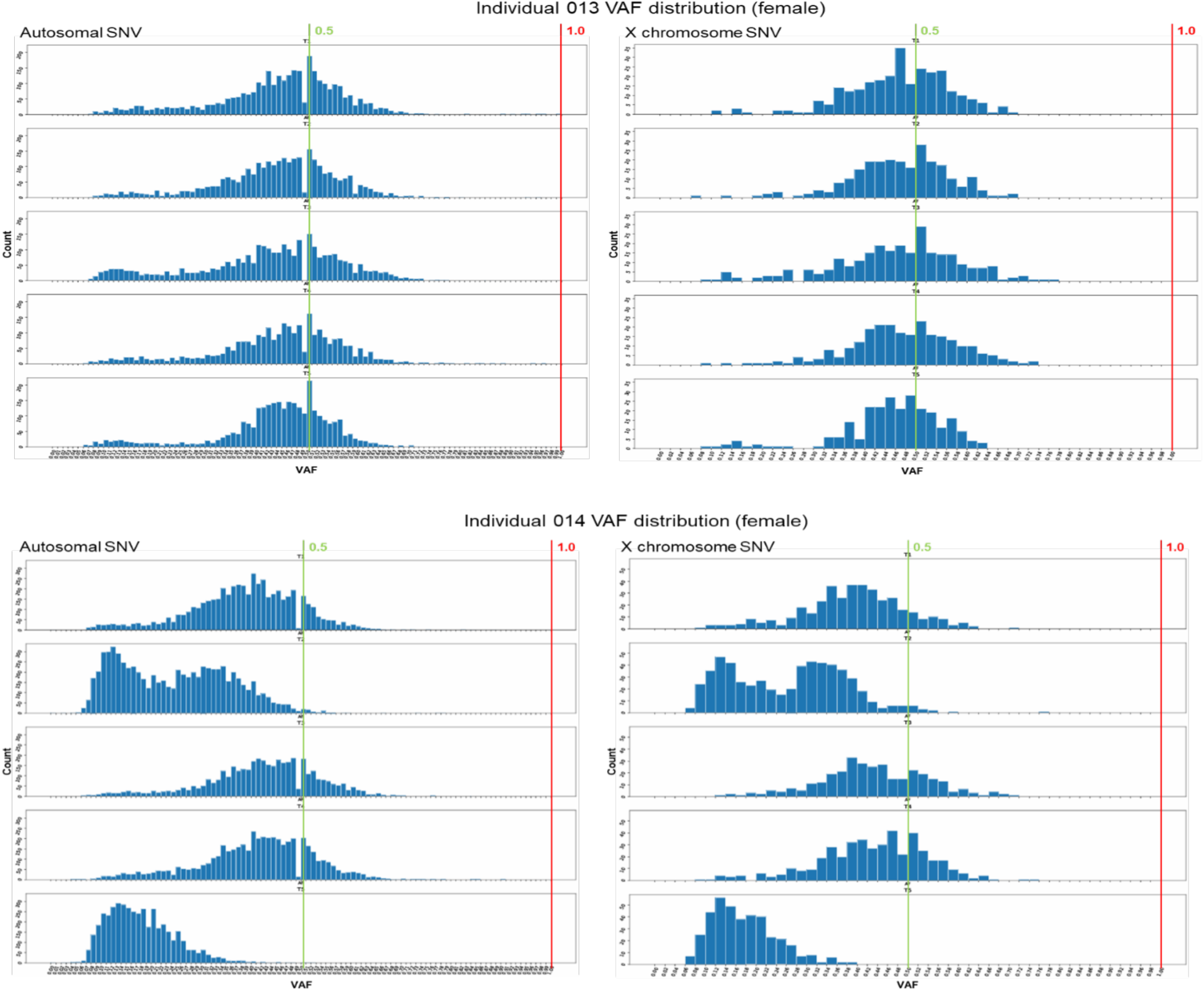
Autosomal and X chromosome single nucleotide variant VAF distribution profile for 71 single human colon crypts. Autosomal SNV calls with a total read count ≥15 from our 2Caller pipeline (common calls from Mutect2 and Strelka) are included in the autosomal SNV analysis. VAF is calculated by dividing the alternate (mutant) allele read count by the total read count. VAF distribution profile is generated for each of the 5 or 6 crypts from 14 individuals. The 14 individuals and their gender designations are indicated with a label on the top of each group of plots with a crypt number marked on the top center (T1 through T6, immediately to the left of the vertical green line) of each panel. The VAF distribution plot for autosomal SNVs is displayed on the left side. The VAF distribution plot for the X chromosome SNVs is displayed on the right side. The Y-axis represents the variant count in each bin of 0.01 displayed from 0 to 1 as VAF on the X-axis for the autosomal VAF distribution plot. The Y-axis represents the variant count in each bin of 0.02 displayed from 0 to 1 as VAF on the X-axis for the X chromosomal VAF distribution plot. VAF 0.5, and 1.0 are marked by the green and the red lines, respectively, as indicated.

**Supplementary Figure S2.**
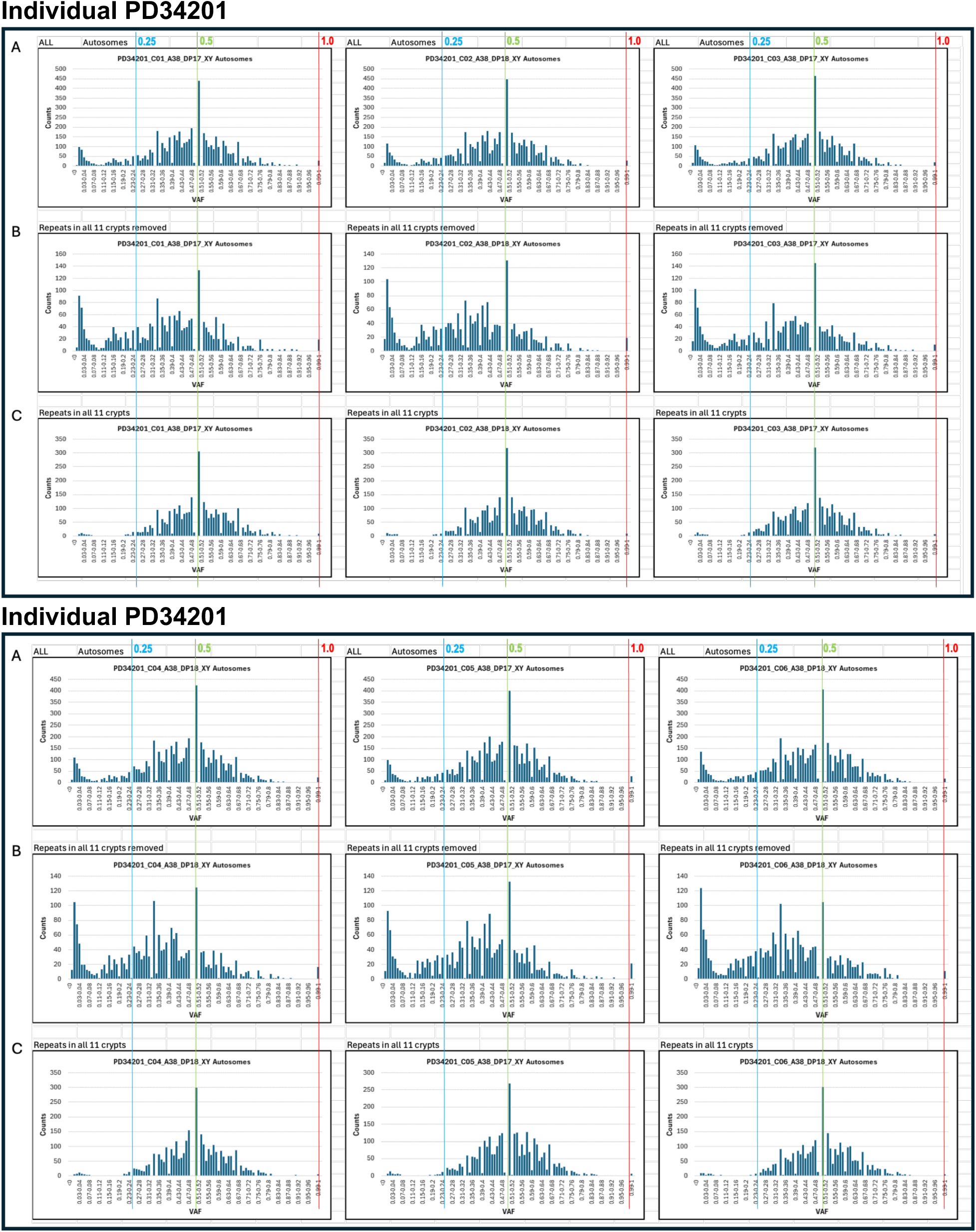
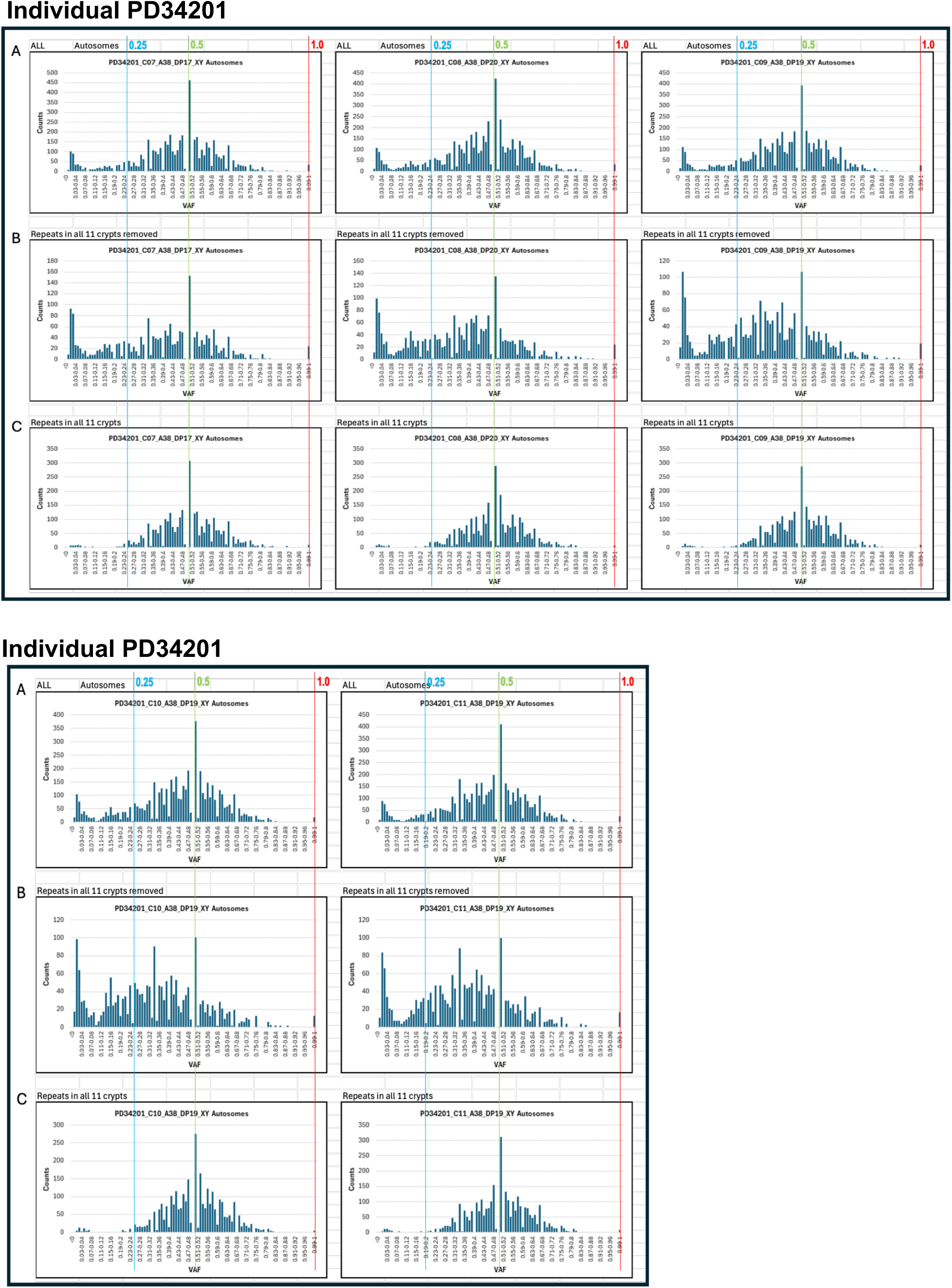
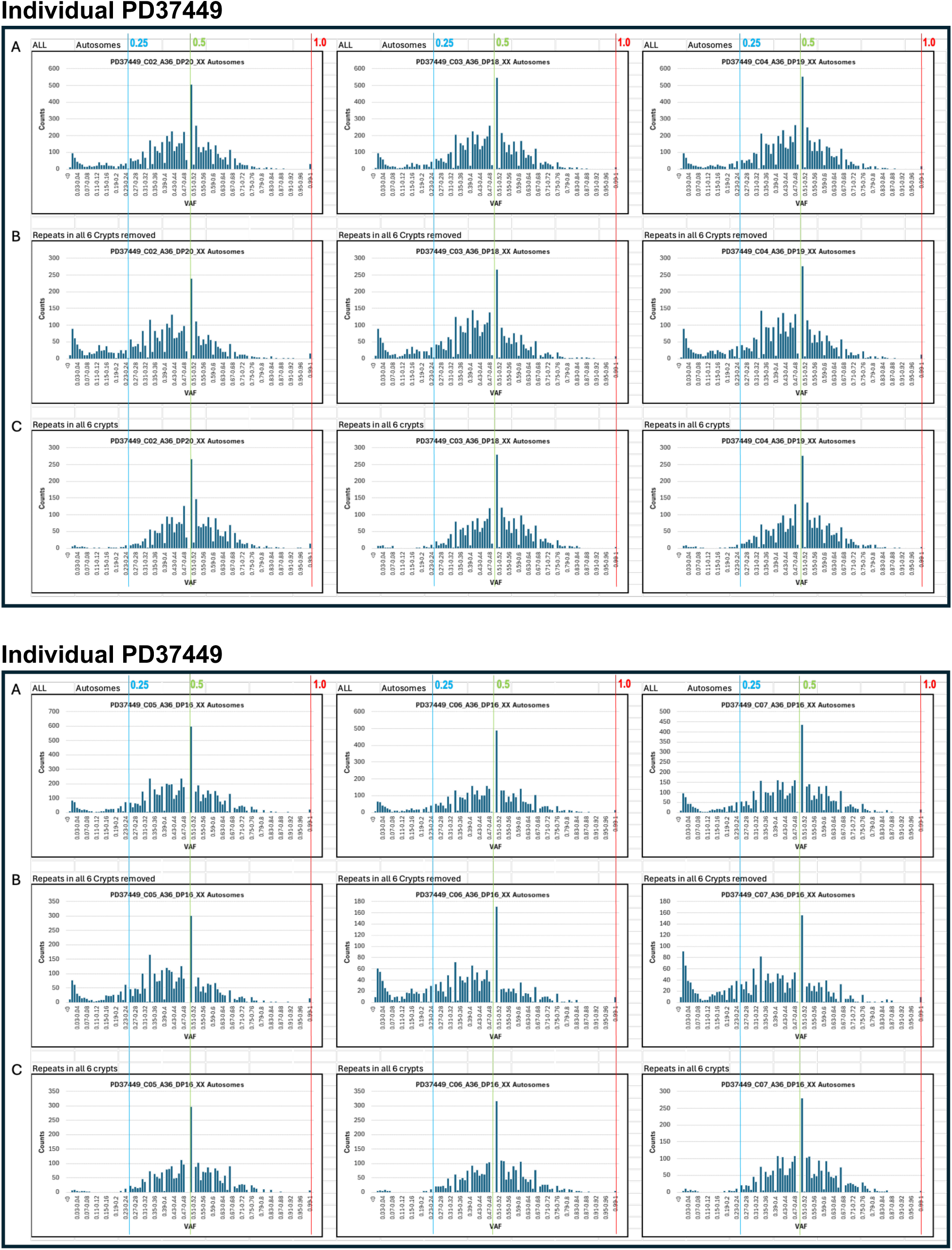
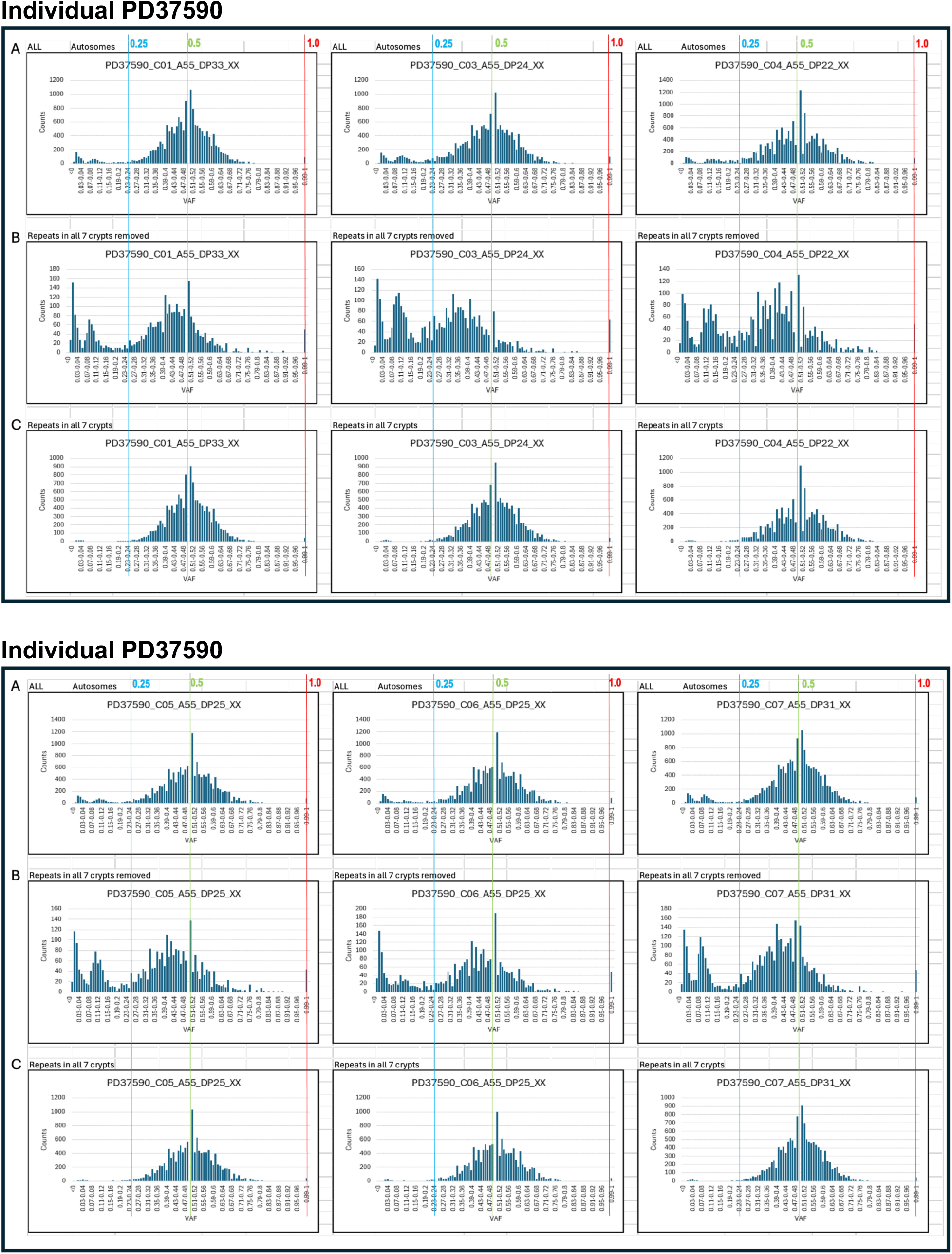
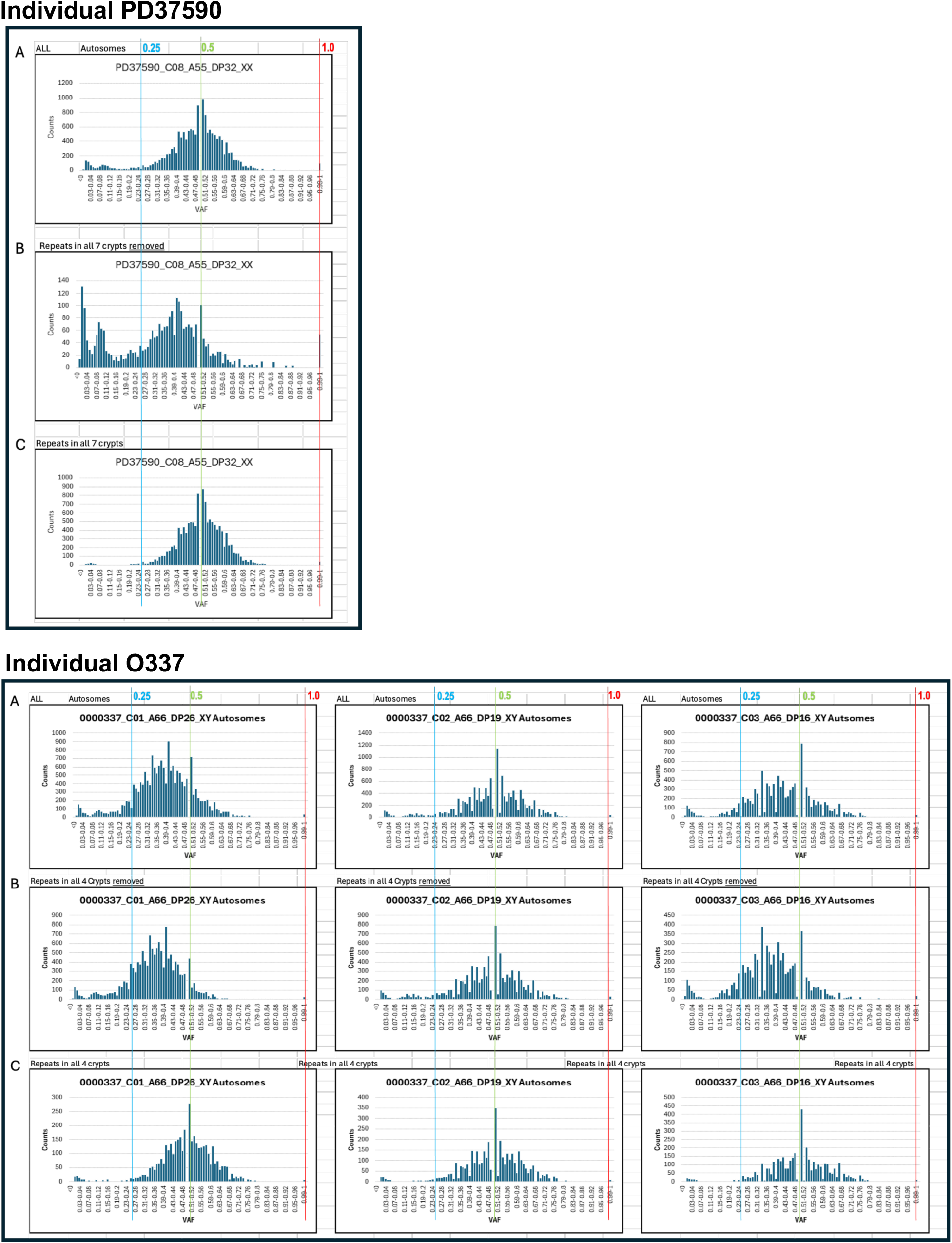
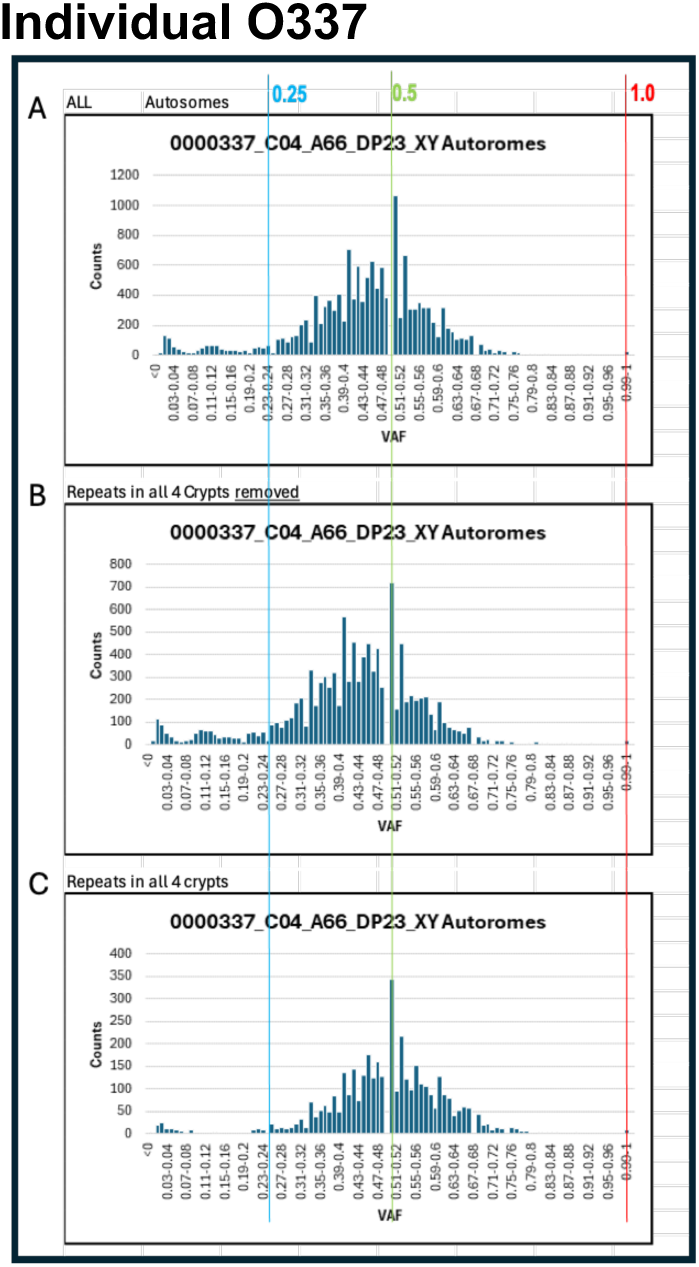
VAF distribution profile of 28 crypts reanalyzed from four individuals in the Sanger study. Each column (consists of A, B, and C plots) represents a single crypt with the patient ID, crypt ID, patient age, median read depth of the sequencing data, and sex chromosome designation displayed with an underscore in between as indicated at the top of the profile. There are a total of 11 crypts from PD34201, a total of 6 crypts from PD37449, a total of 7 crypts from PD37590, and a total of 4 crypts from O337. **Row A)** is the VAF distribution profile of all the autosomal SNV with all the dbSNP variants removed as the first step to reduce germline variant contamination. **Row B)** shows the VAF distribution profile of autosomal variants after the second germline variant contamination reduction step of removing variants shared by all the crypts analyzed from the same individual (repeats in all crypts of the same individual removed). The VAF distribution profiles shown in row B is the best approximation of somatic variants in the Sanger study, in which a matched germline control for each individual is absent (see Supplementary Methods for details). **Row C)** shows the VAF distribution profile of the removed shared variants (repeats in all crypts of the same individual) from the second step of the germline variant contamination reduction. The binomial distribution peaks at VAF 0.5 indicates the germline nature of these shared variants. Deviation of the profile in row B from the profile in row C for the same crypt indicates the multi-lineage nature of the crypt, as shown in the three representative examples in Fig. 3, after germline variants are reduced in the dataset. When nearly identical profiles in row B and row C are observed, as shown in the single representative example in Fig. 3, it can be concluded that the crypt is of single stem cell lineage. Although germline variant contamination cannot be completely removed, the shifted or the extra distribution peak is evident in row B profiles of nearly all 28 crypts. VAF 0.25, 0.5, and 1.0 are marked by the blue, green, and red lines, respectively as indicated. The Y-axis represents the count in each bin of 0.01 displayed from 0 to 1 as the variant allele frequency on the X-axis.

## SUPPLEMENTARY METHODS

### Reanalysis of previously published Sanger Institute low-depth colon crypt WGS data

A conclusion that crypts were derived from a single ancestral stem cell was presented in a previous study (somatic mutations and mutational signatures section of [1]). This conclusion was based on a median VAF distribution plot of 571 microdissected human colon crypts (Extended data Fig. 1d in [1]). In our study, the median VAF of single lineage and multi-lineage crypts in adults showed clear overlap (Figure SM1 below). Therefore, we consider that the median VAF distribution of the 571 crypts presented in the previously published study is insufficient for clonality inference.

**Figure SM1.**
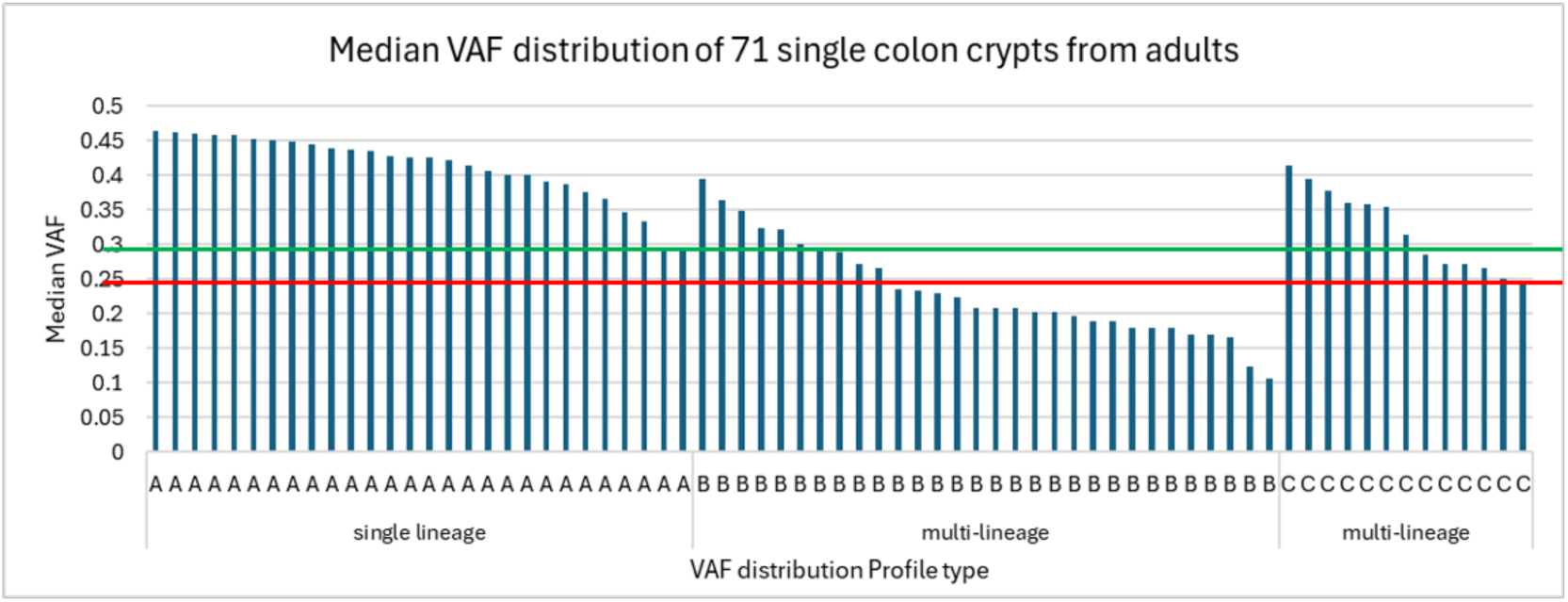
Median VAF distribution of 71 single colon crypts from adults >20 years old. The median VAF of each of the 71 crypts is plotted and grouped by the VAF distribution profile category of each crypt. Profile A crypt is of single stem cell lineage, Profile B crypt is formed by stem cells from multiple lineage without sharing any variants, and Profile C crypt is formed by stem cells branched off from a common progenitor over a long period of time and should be considered as multi-lineage. The green horizontal line marks the lowest median VAF in crypts of Profile A, and the red horizontal line marks the lowest median VAF in crypts of Profile C.

We sought to obtain the WGS data of the 571 crypts in the previous study to generate the VAF distribution profile of each crypt to clarify the cause of different findings in our WGS study. A total of 2038 CRAM files (with multiple CRAM files per sample) for 440 crypt libraries constructed from 42 individuals were downloaded with a signed data transfer agreement (called Sanger study from this point). Although multiple analyses presented in the publication were based on a set of 571 crypts or a set of 445 crypts [1], sequence data for 131 of these crypts and information for 126 of these crypts are unavailable.

Among the 445 crypts in the Sanger study reported in their publication, 276 (62%) have a median depth ≥15X, and 28 (6.3%) were reported to have a median depth ≤10X (Supplemental Table 2 in [1]). Low read depth can jeopardize the accuracy of VAF calculations; therefore, analyzing Sanger samples with a median depth >15X to reduce the impact of low read depth on VAF would be more adequate. It is important to note that there is no matched control sample from each individual in the Sanger study, precluding the accurate elimination of germline variant contamination in the crypt variant calls. Multiple crypts from the same individual can facilitate the reduction of germline variant call contamination in the variant calls in the crypts. Based on these criteria, we chose to analyze samples from four individuals, O337 (age 66 male), PD37449 (age 36 female), PD34201 (age 38 male), and PD37590 (age 55 female), with multiple crypts of median depth >15X.

Eighty-six CRAM files were first merged using Samtools-merge accordingly from 28 samples across four individuals and then were processed through Illumina’s DRAGEN Somatic 4.2.4 pipeline in tumor-only mode with the germline tagging enabled. The summary of the analysis of these 28 crypt samples and corresponding information reported in the Sanger study are listed in Supplementary Table S2. Downstream VAF analysis only includes the autosomal SNV PASS calls from the DRAGEN somatic pipeline after the removal of obvious false positive SNV with germline tags and base quality control assessment. Without a matched germline control, variant calls from each crypt will be substantially contaminated by germline variants even after the initial filtering to increase the confidence of the calls. The infiltrating germline variants can lead to an artifactually high VAF peak at 0.5 and an apparent binomial distribution peak at VAF 0.5 that conceals the true VAF peaks of the unique somatic mutations in each crypt lineage. Germline variant contamination and low read depth at the variant site are the most likely reasons for autosomal VAF 1.0 in the analysis. There are a total of 28 autosomal variants with VAF 1.0 among the 307,229 total variants (<0.01%) and an average of 42.13% (s.d. 9.68%) dbSNP variants in the 71 crypts in our study, which has a matched bulk control from each individual for the somatic variant calls (supplementary Table S1). In contrast, the 28 crypts from the Sanger study have a total of 119,251 autosomal variants of VAF 1.0 among 1,103,691 total variants (average 11.98%, s.d. 3.06%) and an average of 77.66% (s.d. 6.25%) dbSNP variants, indicating the high germline variant contamination in the dataset (Supplementary Table S2). Ideally, we would like to achieve a similar level of VAF 1.0 and dbSNP ratio in our attempt to reduce germline variant contamination in the crypts from the Sanger study. We reasoned that the germline mutations are more likely to be known SNVs, and the true somatic mutations are more likely to be novel. We understand that removing all known SNVs, such as dbSNP, can result in the removal of true positive variants. In addition, variants shared by all crypts from the same individual are more likely to be germline variants, and the probability that the shared variant is a germline variant increases as the number of crypts analyzed from the same individual increases. Therefore, the germline variant contamination can be reduced by removing the variants in the dbSNP database and the variants shared by all crypts from the same individual. The impact of different strategies on germline variant contamination reduction is evaluated by the reduction of variants with VAF 1.0 in each strategy (Figure SM2 and Supplementary Table S2).

**Figure SM2.**
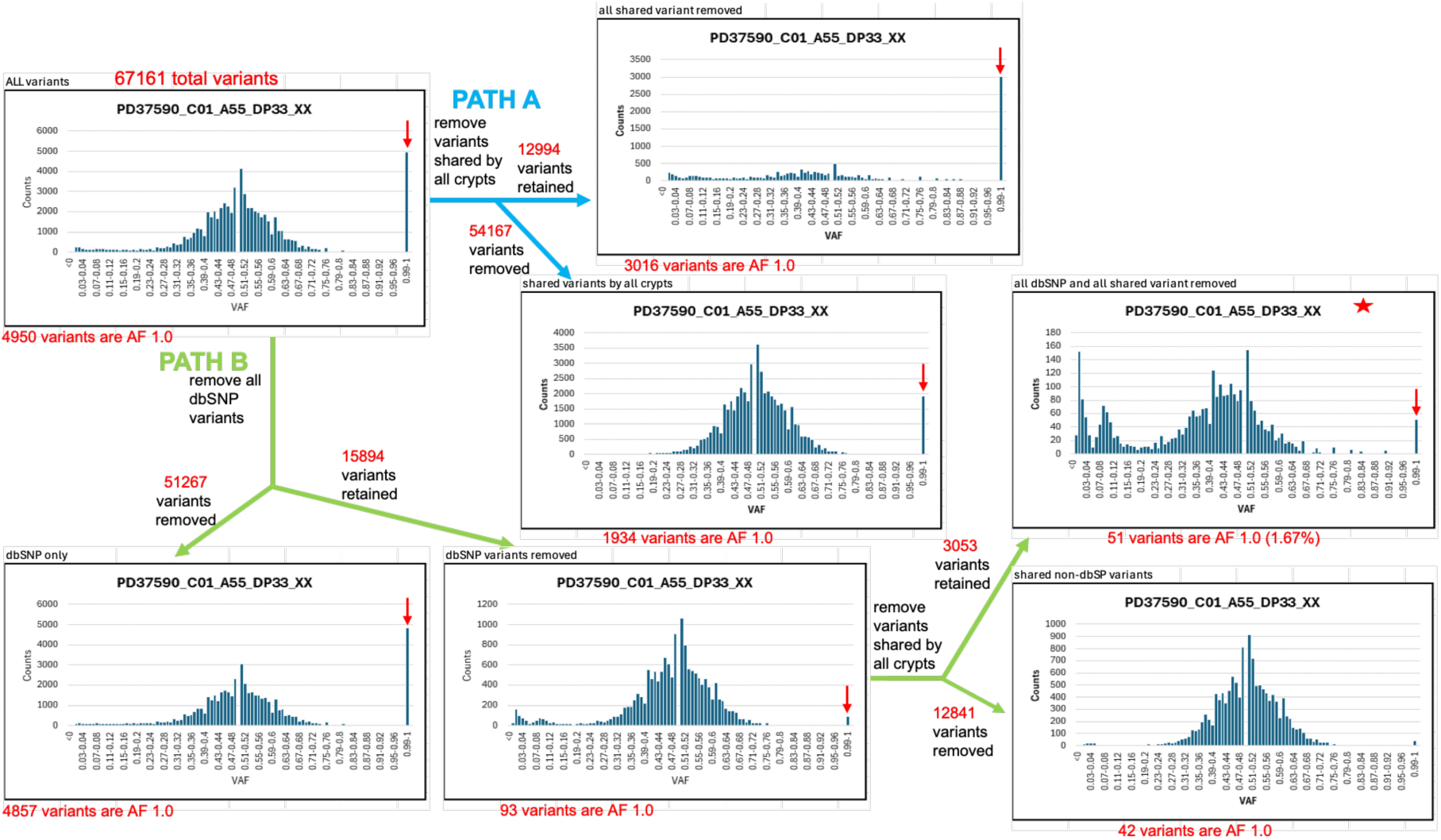
Strategies for germline variant removal. **Path A (in blue)** Variants shared by all 7 crypts from PD3759O are removed. **Path B (in green)** Variants in the dbSNP are removed, followed by the removal of variants shared by all 7 crypts from PD3759O. The number of variants with AF1.0 is used as an indicator for germline variant contamination (indicated under each VAF distribution plot). The red star marks the best outcome for the reduction of germline variants in the crypt. The count of variants removed and retained in each step is indicated at each branch of the path.

The average fraction of dbSNP in the 28 crypts from the Sanger study reduced to 73.09% (s.d. 12.76%) from 77.66% (s.d. 6.25%) and a total of 69,279 variants of AF 1.0 (average 20.42%, s.d. 3.5%) remained after removing all the shared variants in all crypts from the same individual (Figure SM2 path A and Supplementary Table S2). It is clear that removing only the shared variants as in path A is inadequate for reducing germline variant contamination. Therefore, we evaluated whether removing all dbSNP variants would bias the VAF distribution profile of five crypts from an individual representing all three profiles in our WGS study. The VAF distribution profile for each of the three subgroups of SNVs, all autosomal variants, all novel variants, and all dbSNP variants, are generated from five crypts of the same individual in our study (Fig. SM3). VAF histograms of all three subgroups showed highly concordant peak structures (Fig. SM3). This analysis indicates the validity of complete dbSNP removal for reducing germline artifacts without distorting VAF distributions.

**Figure SM3.**
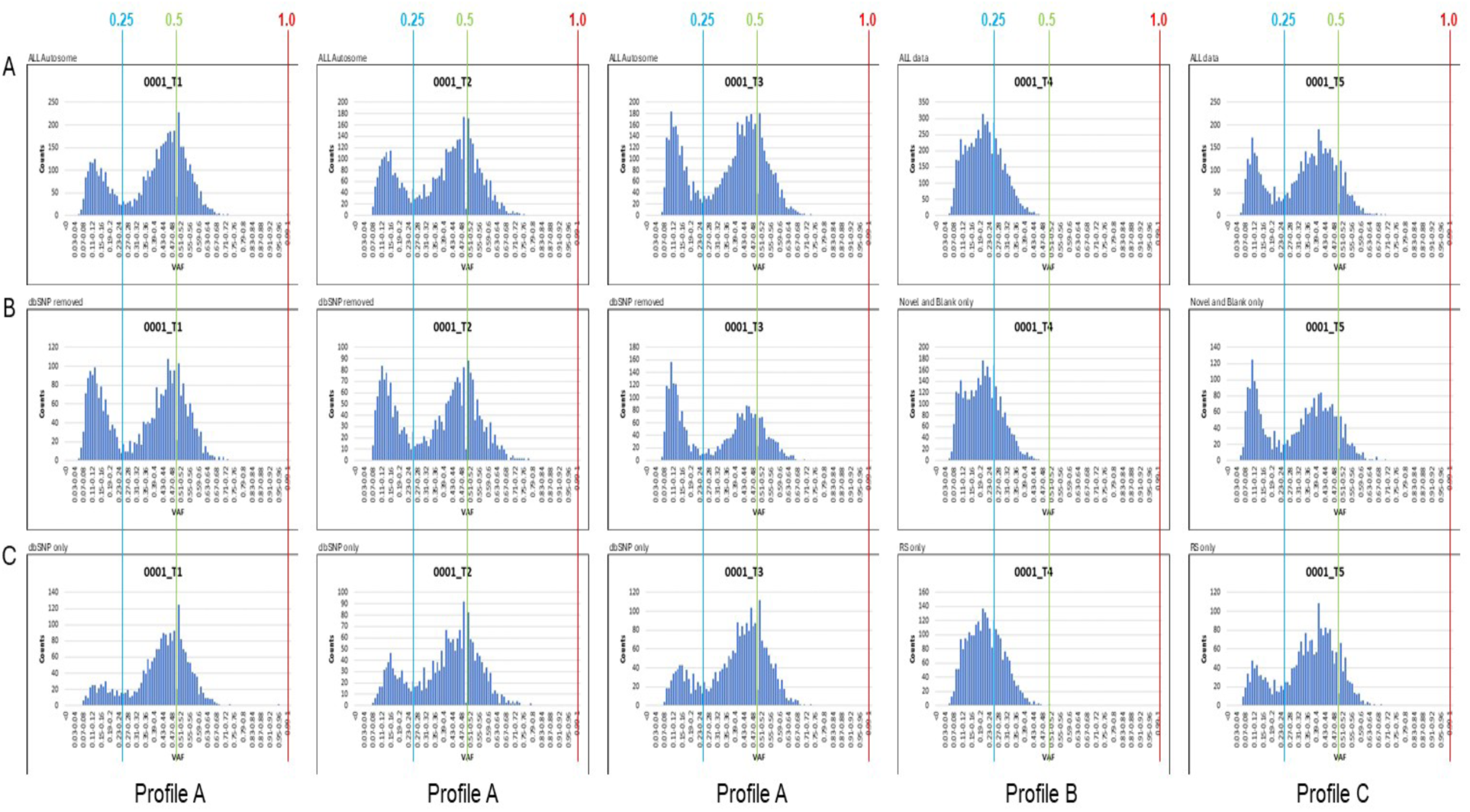
VAF distribution profile of sub-group of variants. Row **A)** VAF distribution profile of all the autosomal SNV with read depth ≥15 in five single colon crypts from the same individual. Row **B)** VAF distribution profile of variants after removal of dbSNP from A) in the same five crypts. Row **C)** VAF distribution profile of dbSNP variants only in the same five crypts. VAF 0.25, 0.5, and 2.0 are marked by blue, green, and red line, respectively, as indicated.

Based on the initial reasoning and the experimental analysis described above, a two-step germline variant removal strategy is devised (Fig. SM2 path B) to remove SNV calls of variants that are in the dbSNP as the first step, and the shared variants in all the crypts of the same individuals are removed as the second step for reducing germline variant contamination in the Sanger study. It is noteworthy that an unusually high number of variants with super high depth are present in the Sanger study (Supplementary Table S2, t_depth >2.5X med_depth column). Variants with super high depth likely represent artifacts from misaligned reads in repetitive regions, and these false positive calls can be reduced by filtering out variants with a 2-fold higher read depth than the median read depth [2]. While such variants may inflate total counts, they typically yield low VAFs and thus do not influence clonality classification based on peak detection between VAF 0.25 and 0.5. Therefore, we retained these variants in the VAF profile analysis

